# White Matter Hyperintensities and Cognitive Decline in de Novo Parkinson’s Disease Patients

**DOI:** 10.1101/230896

**Authors:** Mahsa Dadar, Yashar Zeighami, Yvonne Yau, Seyed-Mohammad Fereshtehnejad, Josefina Maranzano, Ronald B. Postuma, Alain Dagher, D. Louis Collins

**Affiliations:** Montreal Neurological Institute, McGill University, Montreal, Quebec, Canada.

**Author notes:** Corresponding Author Information: D. Louis Collins, Magnetic Resonance Imaging (MRI), Montreal Neurological Institute, 3801 University Street, Room WB315, Montréal, QC, H3A 2B4.

**Keywords:** Parkinson’s disease, white matter hyperintensities, magnetic resonance imaging, cognitive decline, *de Novo* patients

## Abstract

**Objective:** White Matter Hyperintensities (WMHs) are associated with cognitive decline in normative aging and Alzheimer’s disease. However, the pathogenesis of cognitive decline in Parkinson’s disease (PD) is not directly related to vascular causes, and therefore the role of WMHs in PD remains unclear. If WMH has a higher impact on cognitive decline in PD, vascular pathology should be assessed and treated with a higher priority in this population. Here we investigate whether WMH leads to increased cognitive decline in PD, and if these effects relate to cortical thinning

**Methods:** To investigate the role of WMHs in PD, it is essential to study recently-diagnosed/non-treated patients. *De novo* PD patients and age-matched controls (N_PD_=365,N_Control_=174) with FLAIR/T2-weighted scans at baseline were selected from Parkinson’s Progression Markers Initiative (PPMI). WMHs and cortical thickness were measured to analyse the relationship between baseline WMHs and future cognitive decline (follow-up:4.09±1.14 years) and cortical thinning (follow-up:1.05±0.10 years).

**Results:** High WMH load (WMHL) at baseline in PD was associated with increased cognitive decline, significantly more than i) PDs with low WMHL and ii) controls with high WMHL. Furthermore, PD patients with higher baseline WMHL showed more cortical thinning in right frontal lobe than subjects with low WMHL. Cortical thinning of this region also predicted decline in performance on a cognitive test.

**Interpretation:** Presence of WMHs in *de novo* PD patients predicts greater future cognitive decline and cortical thinning than in normal aging. Recognizing WMHs as a potential predictor of cognitive deficit in PD provides an opportunity for timely interventions.

## Introduction

While Parkinson’s disease (PD) is typically characterized by motor symptoms, cognitive deficits occur in approximately 15% of patients in early drug-naïve stages^1^. Two decades after disease onset, this prevalence increases to over 80%^2^. Early mild cognitive impairment (MCI) is a strong predictor of later development of dementia^3,4^, which is a key determinant of mortality and poorer quality of life in PD^5^. Cognitive impairment in PD is related to subcortical dysfunction in early stages, followed by cortical α-synuclein pathology and loss of neurotransmitters. However, it remains unclear to what degree white matter changes, historically described as leukoaraiosis^6^ which are major signs of small-vessel disease (SVD)^7,8^ may contribute to cognitive dysfunction in PD.

White matter hyperintensities (WMHs) or leukoaraiosis are areas of increased signal in T2-weighted and FLAIR structural MRI. The neuropathologic correlates of WMHs are varied: loss of axons and glial cells, myelin rarefaction, spongiosis, perivascular demyelination, gliosis, subependymal glial accumulation and loss of the ependymal lining^8^. Despite the various findings, consensus exists regarding the association of WMHs and SVD^9^. The term SVD is mainly related to two etiologies: 1) age-related vascular disease, also referred as arteriolosclerosis, or vascular-risk-factor related SVD^10,11^, and 2) cerebral amyloid angiopathy^12^. These two play a crucial role in stroke, dementia and aging, and could also be relevant in PD. Therefore, early detection of WMHs and treatment of cardiovascular risk factors could have a positive impact on cognitive decline in PD^13–16^. In AD, WMHs have been extensively studied and strongly predict rapid cognitive decline in individuals with MCI^17,18^. In PD, the pathogenic role of vascular risk factors is less clear^5^ and results have been contradictory^16^. The WMHs might cause cognitive decline independent of PD, or the synergy between the two mechanisms may accelerate cognitive impairment^16^. Alternatively, the WMHs might aggravate the pathologic spread of misfolded α-synuclein or amyloid-β proteins. Of the few studies that have investigated WMHs and cognitive decline in PD, most are cross-sectional, include patients that are on dopaminergic medication, and are typically from cohorts that are at later stages of disease^19–21^. Additionally, different groups implement different tests to assess cognition and many do not perform a comprehensive neuropsychological battery.

Capitalizing on the longitudinal assessment of cognitive abilities and imaging biomarkers in the multi-centre cohort of *de novo* PD patients from the Parkinson’s Progression Markers Initiative^22^, we investigated the relationship between WMH burden and: 1) cognitive decline over time and 2) cortical grey matter changes over time (as indexed by cortical thinning) in early stages of PD.

## Methods

### Patients

The Parkinson’s Progression Markers Initiative (PPMI) is a longitudinal multi-site clinical study of *de novo* PD patients and age-matched healthy controls (HC)^22^ (http://www.ppmi-info.org). The study was approved by the institutional review board of all participating sites and written informed consent was obtained from all participants before inclusion in the study. In the present study, we included all subjects that had either FLAIR or T2-weighted MR images at their baseline visit and had follow-up visits for at least one year after the baseline scan (N_PD_=365, N_HC_=174). All subjects were regularly assessed (yearly follow-ups, mean total follow-up period of 4.09±1.14 years) for clinical characteristics (motor, non-motor and neuropsychological performance) by site investigators, including Montreal Cognitive Assessment (MoCA), Hopkins Verbal Learning Test-Revised (HVLT), Benton judgement of line orientation test for visuospatial skills, Letter-Number Sequencing test for verbal working memory, and semantic fluency test to detect cognitive decline (Table1). The executive function score is calculated as the sum of letter number sequencing and semantic fluency scores^23^. To validate the correlation between these two components, we verified their relationship in the PD population (r=0.56, p<0.0001).

**Table 1.** Descriptive statistics for the PPMI subjects enrolled in this study. Data are number of participants in each category (N), percentage of the total population (%), and mean (SD) of key variables. PPMI=Parkinson’s Progression Marker Initiative. FLAIR= Fluid Attenuated Inversion Recovery. MoCA= Montreal Cognitive Assessment Score. HVLT= Hopkins Verbal Learning Test Revised Total Score. Benton= Benton Judgement of Line Orientation Score. WMH= White Matter Hyperintensity.

|  | Control | *De novo* PD |
| --- | --- | --- |
| Participants (N<sub>Total</sub>) | 174 | 365 |
| Female (N) | 57 (33%) | 114 (32%) |
| T1-weighted and FLAIR Scans (N<sub>Baseline</sub>) | 79 (45%) | 167 (46%) |
| T1-weighted and T2-weighted Scans (N<sub>Baseline</sub>) | 95 (55%) | 198 (54%) |
| Follow-up 3T T1-weighted scans (N<sub>Follow-up</sub>) | 55 (32%) | 100 (27%) |
| Age at Baseline (years) | 60.07 ($\pm 11.34$) | 60.51 ($\pm 9.86$) |
| MoCA at Baseline | 28.25 ($\pm 1.12$) | 27.24 ($\pm 2.22$) |
| HVLТ at Baseline | 35.05 ($\pm 6.78$) | 32.01 ($\pm 7.95$) |
| Benton at Baseline | 26.13 ($\pm 4.12$) | 25.60 ($\pm 4.07$) |
| Executive Function at Baseline | 20.94 ($\pm 4.73$) | 22.29 ($\pm 4.58$) |
| WMH Load at Baseline (cm<sup>3</sup>) | 7.66 ($\pm 10.38$) | 6.93 ($\pm 8.03$) |

### Procedures

All MR images were preprocessed using our standard pipeline^24^ in three steps: noise reduction, intensity non-uniformity correction, and intensity normalization. T2-weighted and FLAIR images were linearly co-registered to the T1-weighted images using a 6-parameter rigid registration. The T1-weighted images were first linearly and then nonlinearly registered to the standard template (MNI-ICBM-152). The WMHs were segmented using a previously validated automatic multi-modality segmentation technique in the native space of FLAIR or T2-weighted scans to avoid further blurring caused by resampling of the images^25,26^. This technique uses a set of location and intensity features obtained from a library of manually segmented scans in combination with a random forest classifier to detect the WMHs in new images. The libraries used in this study were obtained from Alzheimer's Disease Neuroimaging Initiative (ADNI) dataset since the T2-weighted and FLAIR sequences for the PPMI images follow the same acquisition protocol as ADNI. The quality of the registrations and segmentations was visually assessed and cases that did not pass this quality control were discarded (n=43). WMH load was defined as the volume (in cm^3^) of all segmented WMH voxels in the standard space, i.e. the WMH volumes were corrected for total intracranial volume (ICV). All MRI processing and segmentation steps were blinded to clinical outcomes.

For voxel-wise analysis of WMHs, the WMH probability maps generated by the segmentation tool were nonlinearly transformed to the template space at 2×2×2 mm^3^ resolution and blurred with a 3D Gaussian kernel with full width at half maximum of 5 mm to compensate for the variability caused by differences in voxel sizes in the native FLAIR and T2-weighted images. Rates of cognitive decline were calculated for subjects that had at least one-year follow-up information as the change of the score per year (N_PD_=365, N_HC_=174), using a linear regression between time and the score values at different time points along with an intercept term.

Only subjects with T1-weighted 3T MRI data at both initial/baseline visit and at a one-year follow-up MRI were included for cortical thickness analysis (*N_Total_*=155, see Table 1). Cortical models were generated using the CIVET 2.1 preprocessing pipeline^27^, registered to MNI-ICBM-152 template, and analyzed using the SurfStat software package (http://www.math.mcgill.ca/keith/surfstat/). Distances between inner and outer cortical surfaces were evaluated to provide a measure of cortical thickness at each vertex. Changes in cortical thickness were calculated by subtracting the values (Δ*t* = *t*_1_−*t*_2_) at the one-year follow-up (t2) from the baseline (t1). The average time between the baseline and follow-up visits was 1.05±0.11 and 1.05±0.09 years for the PD and control subjects, respectively.

### Statistical Analysis

We tested two major hypotheses: (1) greater WMH load will lead to more extensive and faster decline in cognition of the PD patients (2) patients with a higher WMH load (WMHL) will show more cortical thinning in their follow-up visit after one year.

Survival analysis was used to investigate the relationship between WMH burden and decline in cognition. It has been previously shown that a threshold of WMHs should be present before cognitive deficits are observed^28,29^. The question of interest was whether there is a significant difference between the *cognitive* survival curves of subjects (normal controls and PD patients) with low versus high WMHL. The threshold for differentiating between high and low WMHL was set at 5 cm^3^ (median value, 0.7% of WM volume, 0.27% of brain volume). Similar to previous studies^30–33^, a stable 2-point drop in MoCA (a drop that persists over the follow-up visits) was considered as the terminal event in the survival analysis and the time from baseline MoCA measurement to the visit where the 2-points drop was detected was considered as survival time. This was consistent with recommendations from our in-house clinical consultation. Drop in MoCA was selected as the main terminal event since MoCA has been previously validated as a sensitive measure for detecting and monitoring cognitive change over time^34^ in general and MCI or dementia in PD specifically^35^. Robustness of the results was verified for a WMHL threshold of 10 cm^3^ and 1 to 4 point drops in MoCA. For survival analysis, the survdiff function from R package *survival* was used (ftp://centos.ustc.edu.cn/CRAN/web/packages/survival/survival.pdf). The function implements the two-sample G ^ρ^ statistics family of Harrington and Fleming, with weights on each event (2-point drop in MoCA) of *S*(*t*)^ρ^, where *S*(*t*) is the Kaplan-Meier estimate of survival, i.e. the probability that a subject survives longer than time *t*^36^.

Furthermore, Longitudinal mixed-effects models were used to assess the association of WMHs with changes in cognition. MoCA, Benton, HVLT, and executive function scores were used as measures of cognition (dependent variables). The log-transformed WMH loads and age at each timepoint were used as continuous predictors for either PD or control cohorts. All continuous variables were z-scored prior to the analysis. All models contained first order interactions with age. Subject and contrast used for segmentation (T2-weighted versus FLAIR) were considered as categorical random effects in all the models. Models were fitted using fitlme in MATLAB version R2015b.

Differences in cortical thickness between high and low WMHL classes [(highWMHL_t1_-highWMHL_t2_)-(lowWMHL_t1_-lowWMHL_t2_)] were analyzed statistically based on Gaussian random field theory with a threshold of *p*<0.05^37^. Similar to the survival analysis, the threshold for differentiating between high and low WMHL was 5 cm^3^. Observed differences in cortical thickness were then correlated to cognitive measures using Pearson partial correlations correcting for age.

## Results

### Baseline WMH Load as a Predictor of Longitudinal Cognition

#### Survival Analysis

Baseline WMH loads were not significantly different in control and PD populations (p>0.05). Controlling for age, the rate of decline in MoCA score was significantly correlated with baseline WMH load (r=-0.145, p=0.007) in the PD cohort, but not in controls (r=0.045, p=0.577). Figure 1 shows the Kaplan-Meier plot for the survival analysis for progression decline in MoCA. The 4-year survival rate (i.e. rate of patients maintaining MoCA stability) for the low and high WMHL groups were estimated as 63% (95 CI=0.55-0.70) and 37% (95 CI=0.29-0.45) in PDs and 65% (95 CI=0.52-0.75) and 56% (95 CI=0.45-0.67) in controls, respectively (N_PD_-low=186, N_PD-High_=174, N_HC_-Low=79, N_HC-High_=83). In PD, the high WMHL cohort experienced a significantly lower survival rate than the low WMHL cohort (*χ*^2^=30.9, p<0.00001, hazard ratio= 2.42). There was no high vs low difference in controls (*χ*^2^=2.5, p=0.11, hazard ratio= 1.52). Furthermore, PD patients showed significantly lower survival rate compared to controls in the high WMHL group (*χ*^2^=6.7, p=0.009, hazard ratio=1.58) while the survival rate was not significantly different between two groups in low WMHL group (*χ*^2^=0.1, p = 0. 8, hazard ratio=1.0). Similar results were obtained with a threshold of 10 cm^3^ and 1-4 point drops in MoCA, suggesting that WMH load-based dichotomization is sensitive to a range in the cognitive decline as measured by MoCA.

**Fig. 1.**
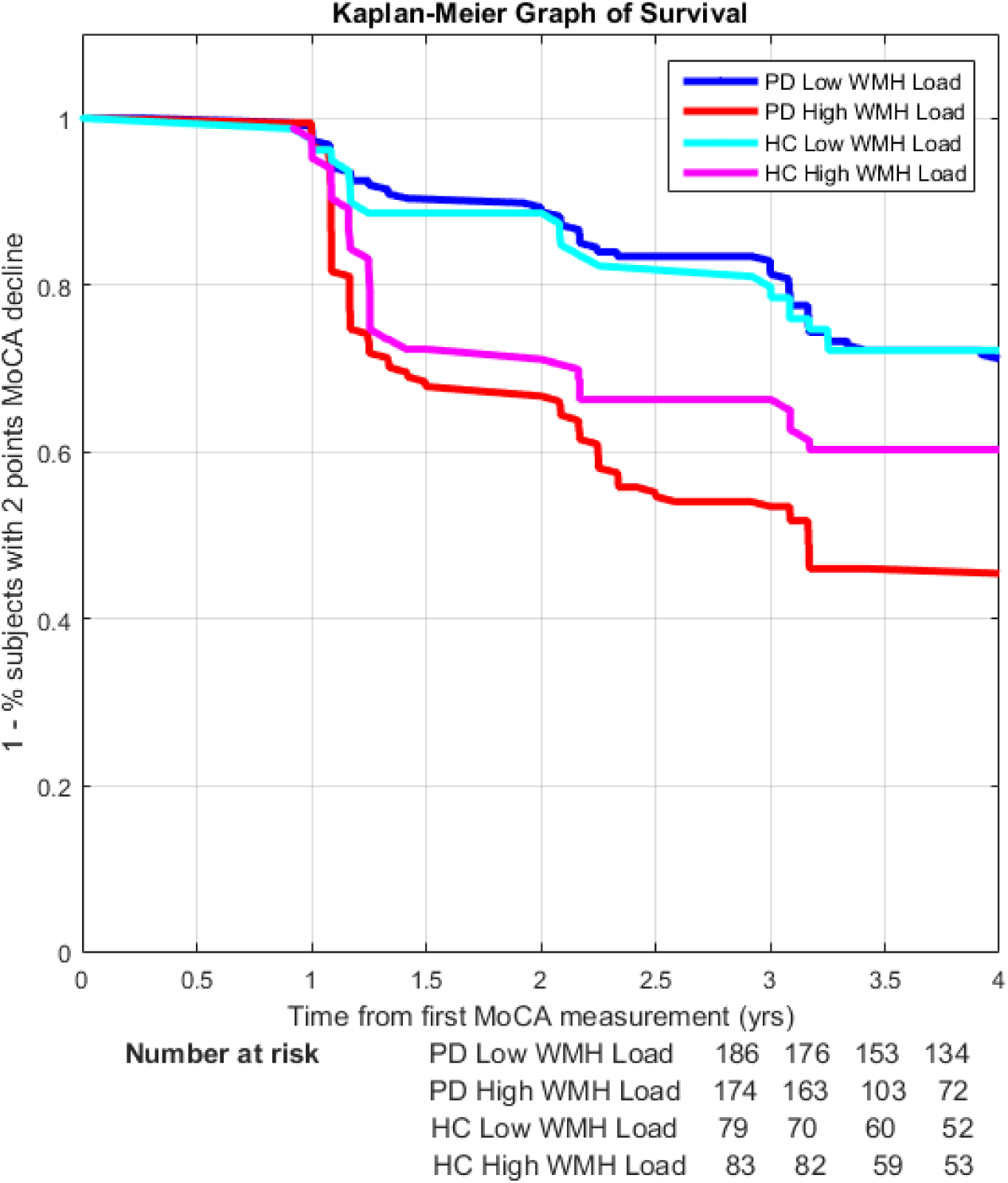
Kaplan-Meier graph of survival showing the survival curves of control and PD patients with low versus high WMH loads demonstrating the compounded affect of PD and WMH load. A 2-point drop in MoCA was considered as the survival event and the time from baseline MoCA measurement to the visit where the 2-point drop occurred was considered as survival time. HC=Healthy Control. PD=Parkinson’s Disease. MoCA= Montreal Cognitive Assessment Score.

### Mixed-Effects Modelling

The mixed-effects modelling results based on age, baseline WMH, and their interaction (Table 2, Fig. 2) showed a significant negative relationship between MoCA, Benton, HVLT, and Executive function scores and age in both PD and HC cohorts. More importantly, in the PD cohort, there was a significant interaction between Age and baseline WMH load for MoCA, Benton, and HVLT which was not observed in the HC cohort.

**Table 2.**
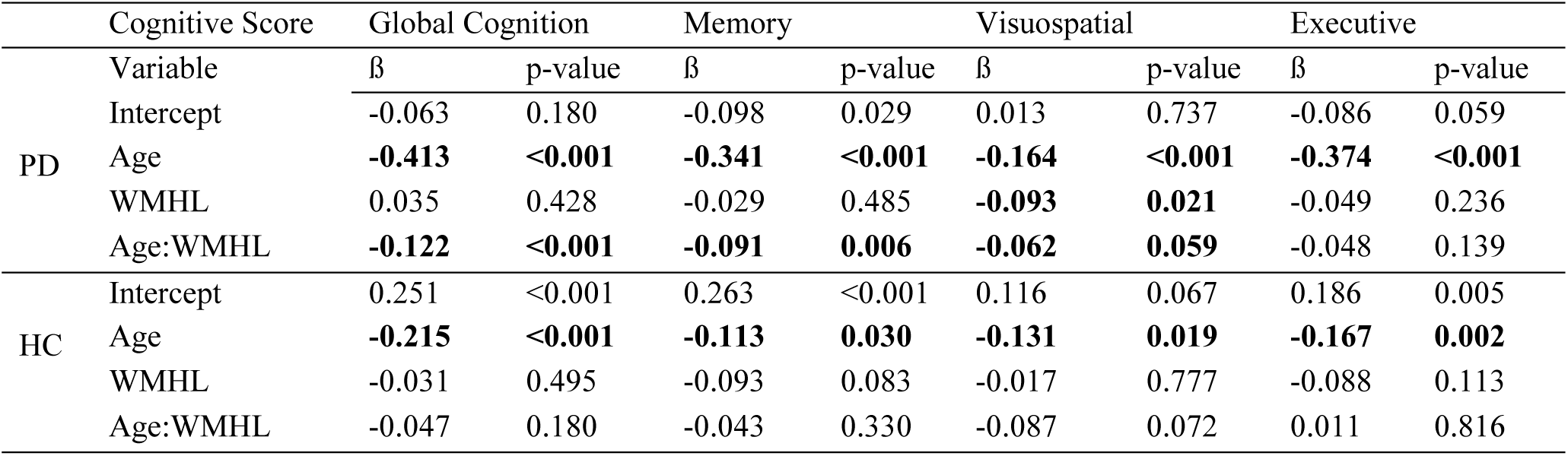
Summary of the mixed effects models of association between baseline WMH Load and cognition in HC and PD cohorts. Entries show the regression coefficients for the listed fixed effect followed by the associated p values. Baseline WMH load was log transformed and z-scored along with age, MoCA, HVLTRT, and Benton scores prior to analysis. WMHL=White Matter Hyperintensity Load. HC= Healthy Control. “:” indicates the interaction between two variables. Global Cognition= Montreal Cognitive Assessment Score (MoCA). Memory= Hopkins Verbal Learning Test Revised Total Score (HVLT). Visuospatial= Benton Judgement of Line Orientation Score. Executive= Executive Function Score (Letter Number Sequencing + Semantic Fluency). HC= Healthy Control. PD= Parkinson’s Disease.

**Fig. 2.**
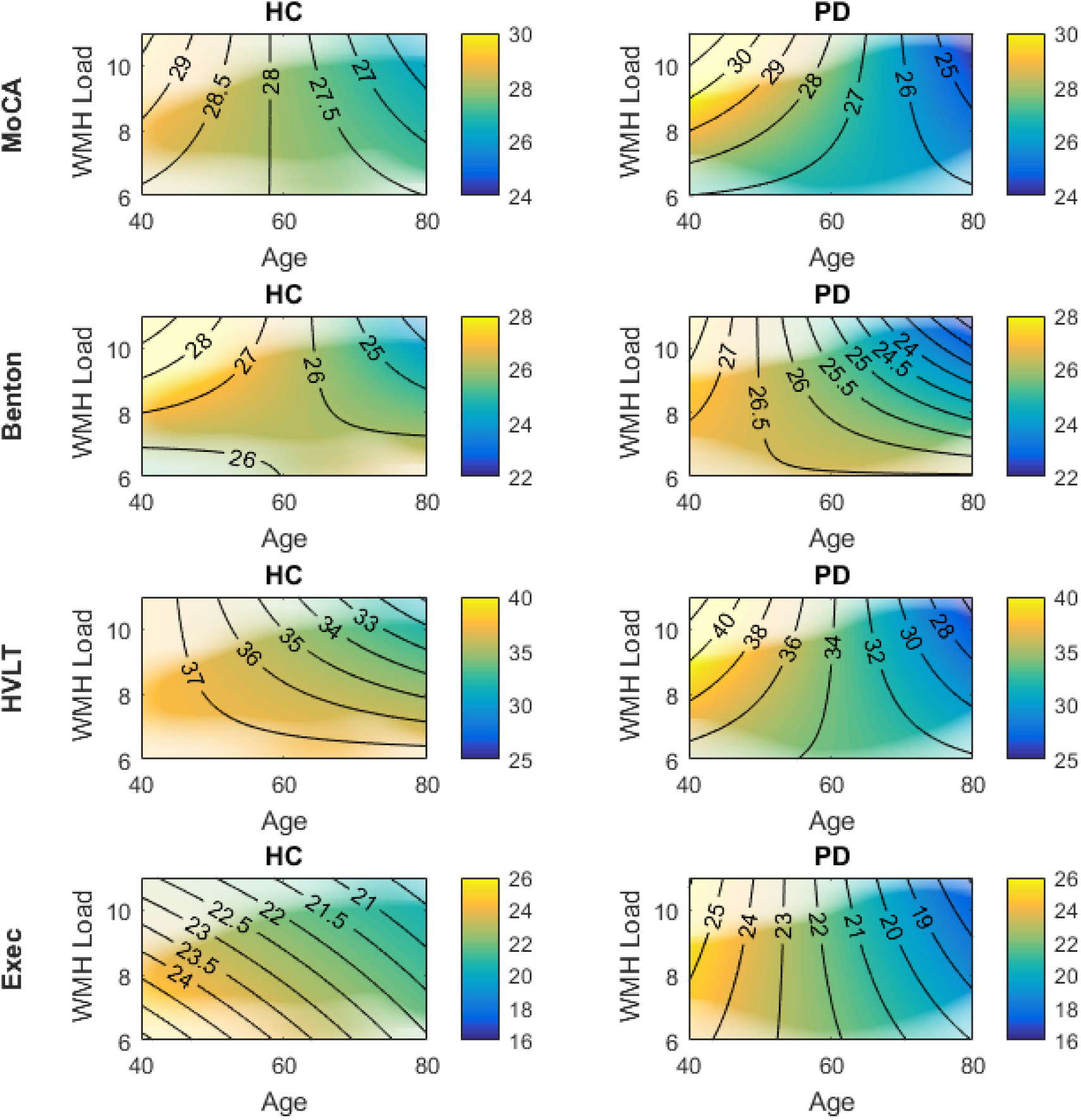
Density plots of longitudinal cognitive changes versus age and log transformed baseline WMH load. The colors indicate predicted cognitive scores by the mixed effects models, with warmer colors representing higher scores, and cooler colors representing lower scores. The transparency in the figures indicates the density of the data, i.e. areas of low transparency indicate regions where there are no subjects and the model is extrapolating (e.g. young subjects with high WMH loads, or old subjects with low WMH loads). The contour lines imply the direction of changes (i.e. horizontal orientation indicates predominance of age effects and vertical orientation indicates predominance of WHM load effects). WMH=White Matter Hyperintensities. HC= Healthy Control. PD= Parkinson’s Disease. MoCA= Montreal Cognitive Assessment Score. HVLTRT= Hopkins Verbal Learning Test Revised Total Score. Benton= Benton Judgement of Line Orientation Score. Exec= Executive Function Score.

### Cortical Thickness

Mean whole-brain cortical thickness decreased significantly among PD patients with both low (t_1_ = 3.3177mm ± 0.0993; t_2_ = 3.3087mm ± 0.1082) and high (t_1_ = 3.2932mm ± 0.0996; t_2_ = 3.2786mm ± 0.0966) WMH at baseline. Among PD patients, baseline WMH load did not correlate with whole-brain cortical thickness at baseline (r=-0.09, *p*>0.05) or at one-year follow-up (r=-0.19, *p*>0.05), but did correlate with cortical thickness change across the one-year period (r=0.26, *p*=0.01). When comparing high and low WMH groups in PD, cortical thinning was greater in the high WMH group with a significant cluster observed in the right frontal lobe (N_Vertices_=1523, resels=7.99, p<0.001) which covers the lateral precentral, superior frontal, and middle frontal gyri (Fig. 3). Cortical thinning of this cluster was not significantly correlated with poorer performance on the HVLT at baseline (r=-0.169, *p*>0.05), but was at one-year follow-up (r=-0.335, *p*<0.001) and with declining performance over the one-year period (*r*=0.196, p<0.05). No significant correlation or vertex/cluster-wise difference was observed in the HC cohort. No significant correlation was observed between MoCA, Benton, and executive function and cortical thickness in PD cohort.

**Fig. 3.**
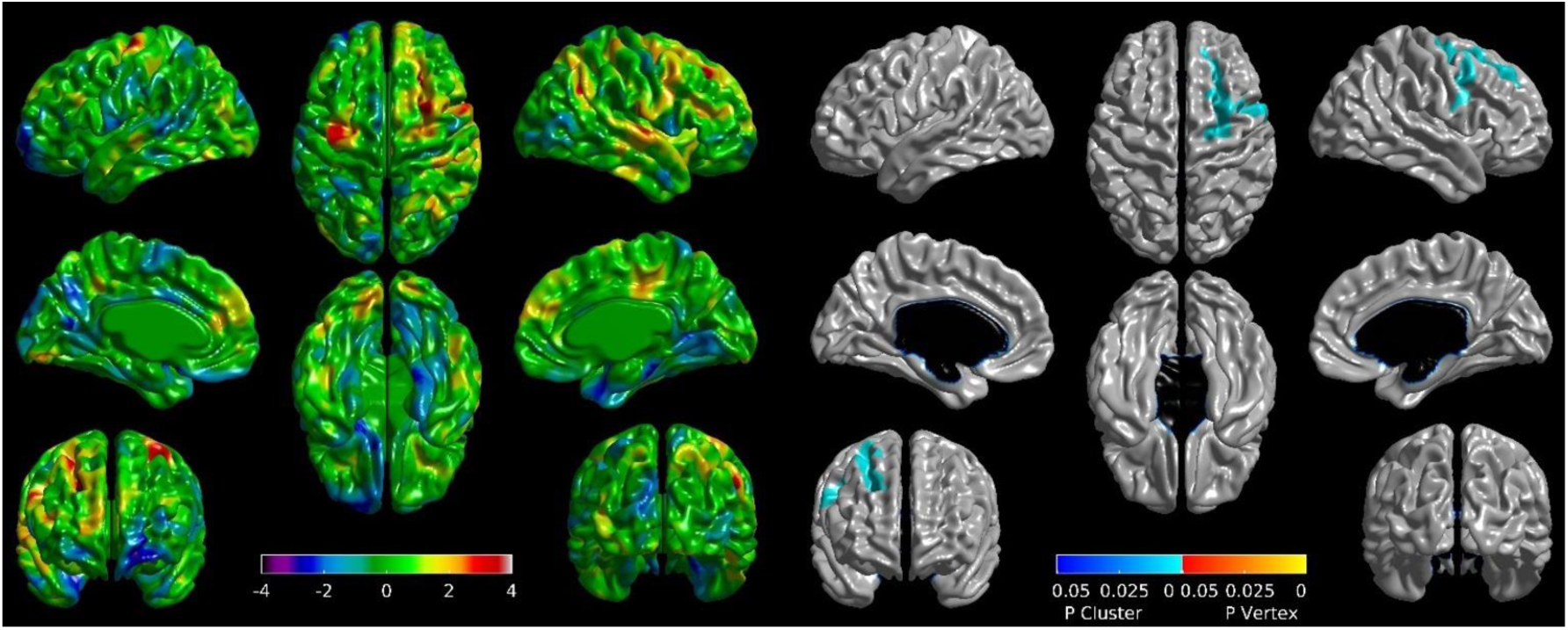
Differences in cortical thickness changes between high and low WMHL cohorts in PD subjects. T-maps (left) and areas of significant cortical thickness decreases (right) covering the precentral, superior frontal, and middle frontal gyri. WMHL= White Matter Hyperintensity Load. PD= Parkinson’s Disease.

### Voxel-wise Analysis

Within the PD cohort, significant voxel-wise correlations were observed between WMH localization maps and the slope of MoCA and Benton scores, corrected for multiple comparisons using false discovery rate (FDR) adjustment and controlled for age and modality (Fig. 4). The significant regions include voxels in all lobes: frontal, temporal, parietal, occipital, and also insular subcortical WM bilaterally. No significant associations were found for the HC cohort. No significant associations were found for HVLT and Executive Function scores in the PD cohort. No significant differences were observed between the baseline voxel-wise WMH maps of PD and HC cohorts after FDR correction.

**Fig. 4.**
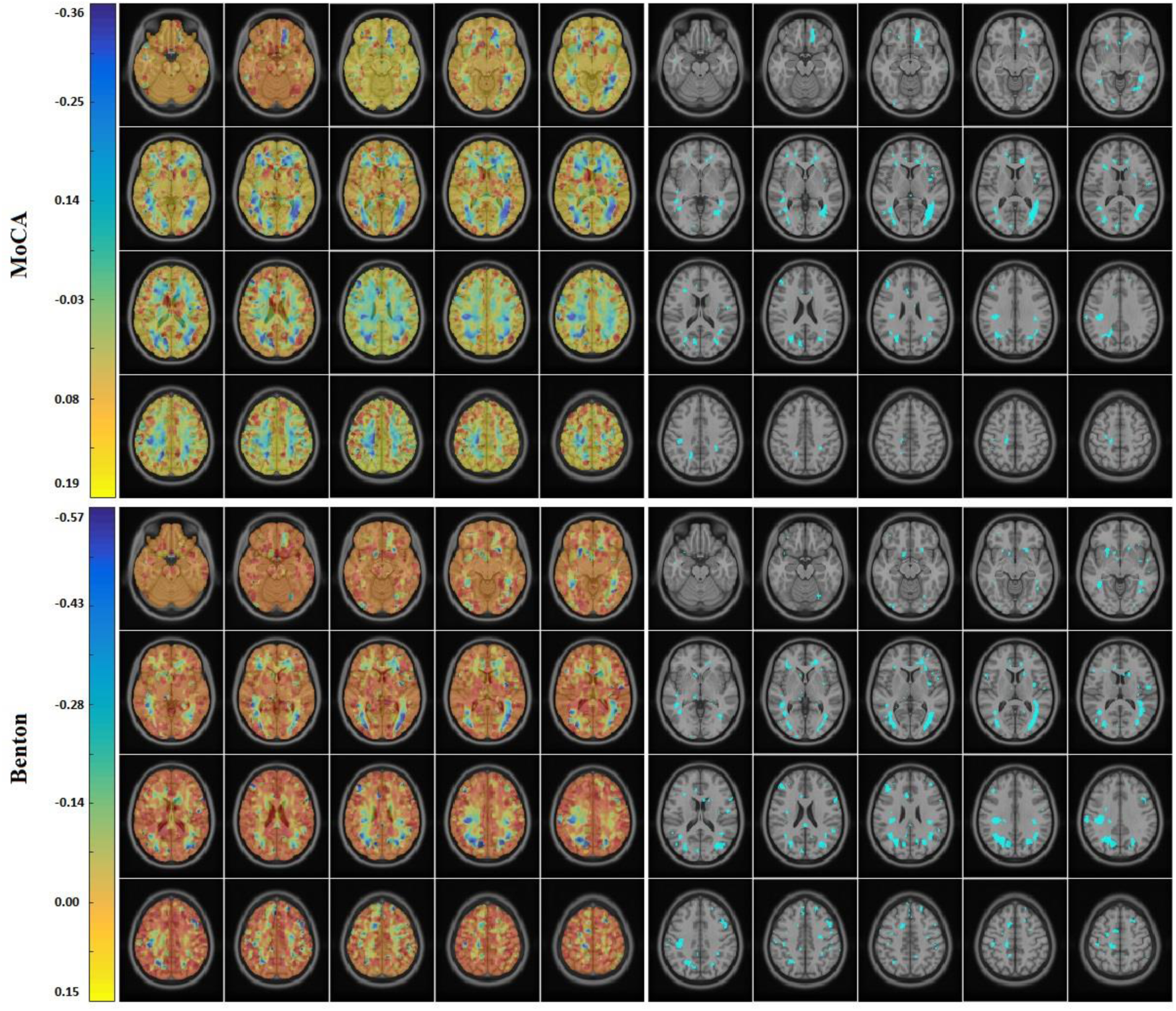
Correlation between WMH location and slope of MoCA (top) and Benton (bottom) score in the PD cohort, controlled for age and modality. Correlation coefficients (left) and thresholded areas of significant correlations after FDR correction. WMH=White Matter Hyperintensity. MoCA= Montreal Cognitive Assessment. PD=Parkinson’s Disease. FDR= False Discovery Rate.

## Discussion

High WMHL PD patients experienced significantly higher decline than i) low WMHL PD patients and ii) high WMHL control subjects. Additionally, WMHL was significantly associated with whole-brain cortical thinning after only one-year follow-up in PD patients, but not in controls. Moreover, PD patients with a high WMHL at baseline showed significant cortical thinning of a frontal cluster compared to those with low WMHL. Taken together, these findings suggest that measures of WMHL in *de novo* PD patients can predict later cognitive decline, even in patients exhibiting no cognitive symptoms at baseline.

As with previous studies^38^, cross-sectional WMHL at baseline in early PD was not significantly associated with baseline cognitive performance. Rather, WMHL at baseline was associated with future cognitive deterioration across multiple cognitive domains including visuospatial, memory, and global cognition corrected for age. This suggests that we can extend previous work on later stages of PD, where WMH burden was significantly associated with conversion to dementia in patients with MCI^39,40^, to the earliest stages of the disease. In line with these findings, post-mortem studies have shown that vascular lesions are common in idiopathic PD (Lewy body disease of the brainstem type)^41^.

MoCA has been validated as a sensitive measure for detecting and monitoring cognitive change over time^34^. Controlling for age, MoCA decline was significantly correlated with baseline WMHL in the PD cohort, but not in controls. Additionally, PD subjects with high WMHLs were more likely to experience a 2-point drop in MoCA than (i) the low WMHL PD and (ii) the high WMHL HC subjects, as evaluated by the survival analysis. The driver for cognitive decline in controls and PD appear to differ in that the former is largely driven by age, while the latter is affected by both advancing age and greater baseline WMH load.

While the literature on PD and WMH is scarce, there has been substantial progress in understanding the relationship between WMHs and cognitive impairment/dementia in AD, especially in the context of amyloid pathology. WMHs associated with vascular risk factors (e.g., hypoperfusion and inflammation) are thought to precede Aβ aggregation. Previous work found significant associations between baseline WMHs and later progression of amyloid load^42^. This further supports the hypothesis of a chain of events; namely WMH impairs clearance of amyloid, which builds up and contributes to cognitive impairment and AD symptoms. While amyloid deposition strongly predicts progression to AD, WMH burden can provide additional independent information to this prediction^43^, suggesting that WMH is not solely related to amyloid pathology, but can directly impact cognitive impairment. Whether a similar interaction between vascular lesions and α-synuclein formation or deposition occurs in PD remains unclear.

WMH burden can also precede irreversible neurological damage as indexed by cortical atrophy. Previous studies have found higher WMHL to be correlated with lower cortical thickness in frontotemporal regions which in turn are related to cognitive decline^44^. Cortical thinning caused by direct or indirect effects of WMHs (tract-specific damage) might lead to cognitive decline and eventually dementia. Cortical thickness might be a sensitive measurement to detect regional grey matter micro-changes that are missed by conventional voxel-based techniques at the earlier stages of the neurodegeneration due to partial volume effect^45,46^. While we observed whole-brain cortical thinning among all PD patients, those with high WMH load showed greater cortical thinning of a frontal cluster, mostly encompassing the right dorsolateral prefrontal cortex (rDLPFC) which was further associated with decline in memory performance in HVLT over the one-year period. This is consistent with previous studies that have found significant associations between rDLPFC and HVLT scores^47,48^. Our results suggest that cortical changes in early PD are potentially moderated by WMH load, and might in turn presage cognitive decline.

Regardless of etiology, prevention and treatment of vascular risk factors associated with WMHs is a promising avenue to slow down cognitive deterioration, especially in *de novo* PD patients who are largely cognitively asymptomatic. The classical and most explored strategy regarding reduction of vascular disease risk and WMHs has been to control hypertension, which subsequently reduces the risk of cognitive deterioration^10,11,49^. In a randomized trial, active lowering of blood pressure was shown to stop or lower the progression of WMHs in patients with cerebrovascular disease over 3 years of follow-up^13^. In the present cohort, we observed an association between WMH load and (systolic-diastolic) blood pressure for both PDs and controls (p<0.001). However, there is also evidence linking WMHs and dementia in PD to orthostatic hypotension, a common occurrence in PD which can be aggravated with anti-hypertensive medication, especially as the disease progresses^50^. This further indicates the need for a tailored blood pressure management in PD patients, while extreme care should be taken to avoid overtreating hypertension. Finally, other small-vessel disease risk factors (some of which have been explored in the context of other pathologies, mainly AD, showing significant correlations with WMHs^15,16^) should be further explored to assess their relevance in WMHs severity and cognitive decline in PD. More importantly, most of these factors are potentially modifiable: percentage of small dense LDL cholesterol, triglycerides level, body mass index, tobacco consumption, type II diabetes, and insulin levels. More studies should focus on assessment of these risk factors in the context of PD and its WMHL.

From a practical standpoint, WMHs can be quantified reliably and non-invasively on large samples and can be measured as a continuous trait, thus providing increased statistical power to detect potential associations^11^. The image processing and WMH segmentation pipelines used in this study have been designed to process data from multi-center studies, are able to control biases due to multi-site MRI scanning (i.e. differences in acquisition parameters), and have been previously applied successfully to a number of multi-site projects^51–53^. The WMH segmentation pipeline has been trained and extensively validated on data from multiple scanners and different acquisition parameters to ensure inter-site and inter-scanner generalizability^26^.

We acknowledge there are limitations to the present study. First, though their differences were accounted for in our analysis, segmentations were based on either T2-weighted or FLAIR images, of which the latter has the better contrast for detecting WMHs. Second, subjects had these scans only at their baseline visit; therefore, we were not able to study the longitudinal changes of WMHs. Future studies investigating WMHs in PD during prodromal and pre-clinical stages are warranted, though there are inherent constraints in recruiting such a cohort. Also, the population under study included relatively cognitively intact individuals (none of the subjects met criteria for dementia), limiting the ability to detect important contributors. Longer follow-ups might further increase the observed differences. One potential confounding factor could be PD medication. However, previous studies have found no significant difference between PD patients on PD medications and PD patients off medications in MoCA and several other cognitive tasks^54^. Similarly, we found no relationship between MoCA and medication in PD patients. Another limitation is that we cannot identify the underlying mechanism. The WMHs might cause cognitive decline independent of PD, however the synergy between the two mechanisms may accelerate the cognitive decline. Alternatively, the WMHs might aggravate the pathologic spread of misfolded α-synuclein proteins in PD. Another possibility is that WMHs in PD may promote amyloid propagation, similar to AD.

In conclusion, our findings suggest that WMH burden is an important predictor of subsequent acceleration in cortical thinning and cognitive decline in early-stage *de novo* PD. Recognizing WMHs as early indicators of cognitive deficit, prior to onset of MCI or dementia, provides an opportunity for timely interventions^22,51^.

## Acknowledgement

We would like to acknowledge funding from *the Famille Louise & André Charron*. Ms. Yau is a Vanier Scholar and receives funding from the Canadian Institute of Health Research. This work was also supported by grants from the Canadian Institutes of Health Research (MOP-111169), les Fonds de Research Santé Québec Pfizer Innovation fund, an NSERC CREATE grant (4140438 - 2012), the Levesque Foundation, the Douglas Hospital Research Centre and Foundation, the Government of Canada, the Canada Fund for Innovation, the Michael J. Fox Foundation and Weston Brain Institute.

Data used in this article were obtained from the Parkinson’s Progression Markers Initiative (PPMI) database (www.ppmi-info.org/data). For up-to-date information on the study, visit www.ppmi-info.org. PPMI is sponsored and partially funded by the Michael J Fox Foundation for Parkinsons Research and funding partners, including AbbVie, Avid Radiopharmaceuticals, Biogen, Bristol-Myers Squibb, Covance, GE Healthcare, Genentech, GlaxoSmithKline (GSK), Eli Lilly and Company, Lundbeck, Merck, Meso Scale Discovery (MSD), Pfizer, Piramal Imaging, Roche, Servier, and UCB (www.ppmi-info.org/fundingpartners). MD and DLC had full access to all the data in the study and takes responsibility for the integrity of the data and the accuracy of the data analysis.

MD, YZ, YY, JM, and AD have no conflicts of interest to report. SMF reports grants from Richard and Edith Strauss Postdoctoral Fellowship, grants from Preston Robb Fellowship, outside the submitted work. RP reports grants and personal fees from Fonds de la Recherche en Sante, grants from Canadian Institute of Health Research, grants from The Parkinson Society of Canada, grants from Weston-Garfield Foundation, grants from Michael J. Fox Foundation, grants from Webster Foundation, personal fees from Biotie, personal fees from Roche/Prothena, personal fees from Teva Neurosciences, personal fees from Novartis Canada, personal fees from Biogen, personal fees from Boehringer Ingelheim, outside the submitted work. DLC reports grants from Canadian Institutes of Health Research, grants from National Science and Engineering Research Council, grants from Canadian foundation for Innovation, other from NeuroRx Inc., other from Truepositive Medical Devices, during the conduct of the study.

## Author Contributions

Mahsa Dadar (MSc): Study concept and design, analysis and interpretation of the data and drafting and revising the manuscript.

Yashar Zeighami (MSc): Study concept and design, analysis and interpretation of the data and revising the manuscript.

Yvonne Yau (MSc): Analysis and interpretation of the data, drafting and revising the manuscript.

Seyed-Mohammad Fereshtehnejad (PhD): Study concept and design, interpretation of the data, drafting and revising the manuscript.

Josefina Maranzano (MD): Interpretation of the data, drafting and revising the manuscript.

Ronald Postuma (MD): Interpretation of the data and revising the manuscript.

Alain Dagher (MD): Study concept and design, interpretation of the data and revising the manuscript.

D. Louis Collins (PhD): Study concept and design, interpretation of the data, drafting and revising the manuscript.

